# Hypoxia Promotes Wound Healing via Dynamical-Mechanical Balance and Adhesion Remodeling

**DOI:** 10.64898/2026.06.11.731571

**Authors:** Zhengduo Wang, Li Tian, Bo Li

## Abstract

Wound healing is a tightly orchestrated physiological process governed by dynamic cell–cell and cell–matrix interactions, yet how hypoxic microenvironments regulate migratory behavior in cells with latent lineage plasticity remains fully elucidated. Here, utilizing human embryonic kidney (HEK293T) and Madin-Darby Canine Kidney (MDCK) cells as a genetically tractable model, we investigate the cellular and molecular mechanisms driving hypoxia-accelerated collective wound repair. Time-lapse live-cell imaging and morphometric analyses reveal that hypoxic exposure significantly accelerates migration, shifts cell cycle dynamics toward the S/G2/M proliferative phases, and induces pronounced morphological spreading. Mechanistically, hypoxia induces a persistent, time-dependent downregulation of the desmosomal cadherin desmoglein-2 (DSG2), thereby weakening intercellular cohesion. Concurrently, the cell-matrix adhesion molecule integrin β3 (ITGB3) exhibits a distinctive biphasic kinetic response—an initial sharp upregulation followed by a sustained decline-which serves to optimize focal adhesion traction and subsequent trailing-edge detachment. Transcriptomic profiling further corroborates these phenotypic transitions, demonstrating a global enrichment of gene networks associated with plasma-membrane adhesion organization, receptor activity, and ion homeostasis that independently mirrors the altered junctional dynamics and accelerated cellular responses. Collectively, our findings uncover a novel cooperative mechanism by which hypoxic stress coordinates cell-cell and cell-matrix adhesion remodeling to facilitate efficient tissue repair, highlighting the valuable utility of plastic cellular models in decoding microenvironmental stress responses.

## Introduction

Wound healing is a highly coordinated, multi-stage biological process that strictly depends on the synergistic actions of cell proliferation, morphological remodeling, and directed collective migration [1]. Following tissue injury, the local microenvironment undergoes rapid and drastic changes, most notably a sharp decline in oxygen tension, which creates a physiological hypoxic microenvironment [2, 3]. This hypoxic milieu acts as a potent physiological signal that activates an evolutionarily conserved cellular stress response pathway centered on the hypoxia-inducible factor (HIF) family of transcription factors [4]. Through both HIF-dependent and HIF-independent mechanisms [5], hypoxia triggers global cellular reprogramming, endowing adherent epithelial and mesenchymal cells with enhanced migratory capacity, metabolic flexibility, and stress resistance—adaptive changes that are essential for efficient wound closure [6-8]. However, despite decades of investigation, the cell-type-specific responses to hypoxia and the precise molecular mechanisms that link oxygen sensing to the dynamic regulation of cell adhesion and motility remain incompletely elucidated.

Human embryonic kidney (HEK293T) cells are a widely used human embryonic kidney-derived cell line that has traditionally been regarded as a stable and genetically tractable tool for molecular biology research [9]. In scratch-wound assays, HEK293T cells have been effectively used to demonstrate how specific scaffolding proteins and macromolecular motor complexes, such as the NAGK–NudC–Lis1 complex, promote directed cell translocation to accelerate gap closure [10]. Concurrently, Madin-Darby Canine Kidney (MDCK) cells represent the gold-standard model for collective epithelial cell migration and barrier morphogenesis due to their distinct apicobasal polarity and robust junctional complexes [11, 12]. Extensive research using wounded MDCK monolayers has established that orderly wound closure requires a sophisticated, dynamic coordination of cell-matrix and cell-cell adhesions, where the remodeling of focal integrins and the spatial redistribution of desmosomal proteins critically transition the epithelium from a quiescent state to a dynamically migrating sheet [13, 14]. However, while the mechanical and signaling profiles of these two individual cell lines have been characterized under normoxic baselines, the precise mechanisms by which hypoxic stress modulates their respective wound healing responses and drives adhesive junctional remodeling remain largely unexplored.

In the present study, we established a hypoxia-exposure experimental model using HEK293T and MDCK cells in a 2D culture system and employed live-cell imaging to capture collective cell behaviors, with the aim of dissecting the cellular and molecular mechanisms underlying hypoxia-accelerated migration. We found that hypoxic conditions (2% O2) markedly enhanced both collective and single-cell motility of HEK293T cells in standard 2D culture, accompanied by cell cycle acceleration (S/G2/M phase transition) and changes in cell spreading morphology. Mechanistically, live-cell imaging captured a persistent, time-dependent downregulation of desmoglein-2 (DSG2) expression under hypoxia, as well as a biphasic expression pattern of integrin β3 (ITGB3), which initially increased and then declined continuously. Transcriptomic profiling further corroborated this migratory phenotype, revealing a comprehensive enrichment of gene networks regulating plasma-membrane adhesion, receptor activities, and intracellular ion homeostasis that independently mirrors the observed junctional remodeling and accelerated cellular responses. Collectively, our findings uncover a novel mechanism by which hypoxia promotes wound-healing migration through the dynamic regulation of cell–cell and cell–matrix adhesion in both HEK293T and MDCK cell systems. Furthermore, this study highlights the indispensable power of time-lapse live-cell imaging in decoding microenvironmental stress responses, which uniquely enabled the real-time tracking of cellular plasticity—such as capturing the precise, biphasic kinetic response of ITGB3 that transiently peaks before declining—thereby revealing how these distinct cell models temporally coordinate their adhesive machinery to facilitate efficient epithelial repair.

## Materials and Methods

### Cell line maintenance and culture

HEK293T (ATCC) cells were grown in DMEM (Adamas life, C8013-500mL) supplemented with 10% heat-inactivated FBS (Gibco). MDCK (ATCC) cells were grown in DMEM/F12 (Adamas life, C8015-500mL) supplemented with 10% heat-inactivated FBS (Gibco). Cells were maintained in an incubator at 37□ □ and 5 % CO2 and passaged every 2 to 3 □days. The mycoplasma testing was conducted weekly on the cell lines using the Mycoplasma Detection Kit (Solea, CA1080). We use Corning Matrigel (356234-5mL) to create a 3D culture environment that assists the tumor cells in forming spheroids. After thawing the Matrigel according to the manufacturer’s recommendations, mix it thoroughly with a pipette to ensure homogeneity. Dilute the Matrigel to a 50% concentration with a serum-free medium. Then, pour the diluted Matrigel on the confocal dish and incubate at 37°C for 15 minutes to allow solidification. Finally, seed the cells on the confocal dish with solidified Matrigel and add complete culture medium.

### Construction of monoclonal genetically modified cell lines

We utilized overexpression lentiviral vectors to add fluorescent signals to the target protein for visualization imaging and of target proteins. The lentiviral vectors used in this study were constructed and synthesized by GeneChem (https://www.genechem.com.cn/about.html). We packaged the synthesized vectors into lentivirus. The GeneChem lentiviral system consists of the GV lentiviral vector series, pHelper 1.0, and pHelper 2.0 plasmids. After mixing the plasmids, we performed transient transfection into HEK293T cells using the E-trans DNA transfection reagent (GeneChem, 10038555). Following gentle mixing, the cells were incubated for 48 hours, and the supernatant was collected. Lentivirus was purified using the GeneChem lentiviral particle purification kit and stored at -80 °C. The day before infection, cells were counted, and approximately 10,000 cells were seeded in a 12-well plate. After an overnight culture, the virus was diluted in serum-free medium and added to the plate to infect the cells at the recommended MOI. After 12 hours, the virus-containing solution was removed, and a complete medium was added for cell culture. After 48 hours, fluorescence intensity was observed under a fluorescence microscope. Corresponding antibiotics were added for cell selection, and monoclonal clones were picked for subsequent experiments. Eight stable cell lines are constructed for this study: HEK293T-LifeAct-H2B to visualize F-actin and H2B, HEK293T-FUCCI to visualize cell cycle, HEK293T-DSG2 to visualize DSG2, HEK293T-ITGB3 to visualize ITGB3, MDCK-LifeAct-H2B to visualize F-actin and H2B, MDCK-FUCCI to visualize cell cycle, MDCK-DSG2 to visualize DSG2 and HE MDCK-ITGB3 to visualize ITGB3.

### Immunostaining Experiments

For the samples in 3D, the culture medium was aspirated, and the samples were washed with PBS. They were then fixed at room temperature with 0.4 % glutaraldehyde (National Medicines, 30092436) for 15 minutes, followed by another wash with PBS. Next, the samples were treated with 0.1 % sodium borohydride (Kemiou, 80115818KMO) at 4 °C for one hour to eliminate the effects of glutaraldehyde on imaging. After washing with PBS again, the samples were permeabilized at room temperature with 0.1 % Triton X-100 (Solarbio, T8200-500ml) for 30 minutes. They were then blocked at room temperature for 2 hours using a solution of 3% BSA (Pumeike, PMK0181-250g) prepared in 0.1 % Triton X-100. The samples were incubated overnight at 4°C in the dark with Alexa Fluor™ 488 phalloidin (Thermo Fisher, A12379) diluted 2000-fold in 3 % BSA for actin. The diluted antibody was incubated with the 3D samples overnight at 4 °C. After washing twice with PBS, an anti-fade mounting medium containing DAPI (Pumei Biological, PMK0272) was added, and the samples were incubated at room temperature for 15 minutes before imaging.

For the samples in 2D, the culture medium was aspirated and washed once with PBS. They were then fixed at room temperature with 4% paraformaldehyde (Pumei Biological, PMK2040) for 15 minutes, followed by another wash with PBS. The samples were permeabilized at room temperature with 0.5% Triton X-100 (Solarbio, T8200-500ml) for 10 minutes and then washed again with PBS. Next, a blocking solution of 3% BSA (Pumei Biological, PMK0181-250g) prepared in PBS was applied for 1 hour at room temperature. The samples were incubated overnight at 4°C in the dark with Alexa Fluor™ 488 phalloidin (Thermo Fisher, A12379) diluted 2000-fold in 3 % BSA for actin.The DSG2 primary antibody (Abcam, catalog no. ab109445) was diluted 500-fold with 3 % BSA. The E-cad primary antibody (Abcam, catalog no. ab11512) was diluted 500-fold with 3 % BSA. On the following day, the antibody was removed, and the samples were washed three times with PBS, each wash lasting 10 minutes. The Goat anti-Mouse or Rabbit IgG (H+L) secondary antibody was then diluted 1000-fold in 3% BSA and incubated with the samples for 2 hours at room temperature in the dark. Finally, the samples were washed three times with PBS, each wash lasting 10 minutes. After washing twice with PBS, an anti-fade mounting medium containing DAPI (Pumei Biological, PMK0272) was added, and the samples were incubated at room temperature for 15 minutes before imaging.

### Live cell imaging

For live imaging, we established a live cell imaging platform by integrating a 3-gas controller (Oko-lab, 3GF-MIXER-HYPOXIA) and a temperature incubator (Oko-lab, H301-MINI) onto a spinning disk confocal microscope (SpinSR, Evident Olympus). The microscope incubators maintain a temperature of 37°C for live cell culture, while the gas controller regulates the flow rates of air, carbon dioxide, and nitrogen to maintain a carbon dioxide concentration of 5% and an oxygen concentration of either 21% or 2%, depending on experimental requirements. Cell samples in a confocal dish (Nest, 801001) were placed on the live cell workstation. Typically, we set the laser intensity to 20% and the exposure time to 200 ms. To capture the dynamics of tumor spheroids, we utilized a laser with a wavelength of 488 nm to image the cytoskeleton and a laser with a wavelength of 561 nm to image the nuclei. To capture cell cycle, the 488 nm laser was used to image nuclei in the S/G2/M phases, while the 561 nm laser was employed to image nuclei in the G1 phase. When imaging 2D samples, at each position, a total 15 µm range along the z-axis was scanned at an interval of 1 µm, resulting in a total number of 15 images for one stack. The super-resolution images are obtained under the corresponding mode of the spinning disk confocal microscopy, using a 60x objective. Images were captured every hour for a total duration of 42 hours, with no significant phototoxicity observed

### Analysis of live cell imaging data

To analyze the cellular morphology, we segmented the cell using CellPose [15, 16], which provided the mask file of the cells. XY coordinates of each point on the cell’s edge are then calculated based on the mask data. Subsequently, using the segmentation data, we employed custom-written MATLAB code to analyze dynamic parameters, including area, perimeter, circularity, movement direction, and size of the cells. Time-series image stacks were imported into ImageJ. The Tracking Plugin Manual Tracking was utilized to obtain the movement trajectories, centroid spatial coordinates, distance, and speed of the tumor spheroids.

### Transcriptome sequencing and analysis

The cells cultured in two dimensions can be lysed directly using Trizol. After culturing cell spheroids under normoxic or hypoxic conditions for 24 hours, the culture medium was aspirated, and TransZol (TransGen, ET101-01-V2) was added directly to the cells. RNA was then extracted and purified using the EasyPure® RNA Kit (TransGen, ER501-01-V2). The purified RNA was sent to Beijing Genomics Institution for library construction and sequencing, yielding sequencing data. We subsequently utilized R packages and Galaxy (https://usegalaxy.cn/) to align the sequencing data with human genomic information. To elucidate patterns and regulatory mechanisms of gene expression, regular analyses have been conducted, including sequence alignment, gene expression quantification, and differential expression analysis. Furthermore, we conducted GO and KEGG enrichment analyses of differentially expressed genes to gain deeper insights into the molecular mechanisms and regulatory networks underlying biological processes.

## Results

### Dimensional Transition from 2D to 3D Microenvironments Triggers a Profound Phenotypic Switch in Epithelial Cells

To delineate the differences in phenotypic switching capacity between HEK293T cells and classical epithelial cells, we established both 3D and 2D culture platforms for HEK293T and MDCK cells in parallel and performed comparative analyses [17]. Immunofluorescence staining revealed strikingly divergent morphogenetic programs: MDCK cells successfully assembled into hollow, polarized tubular structures in 3D matrices, a hallmark of normal epithelial differentiation and lumen formation, whereas HEK293T cells aggregated to form compact, solid tumor-like spheroids, indicating substantial differences in cell–cell adhesion and polarization capacity (**Fig. 1A-B**).

**Fig. 1.**
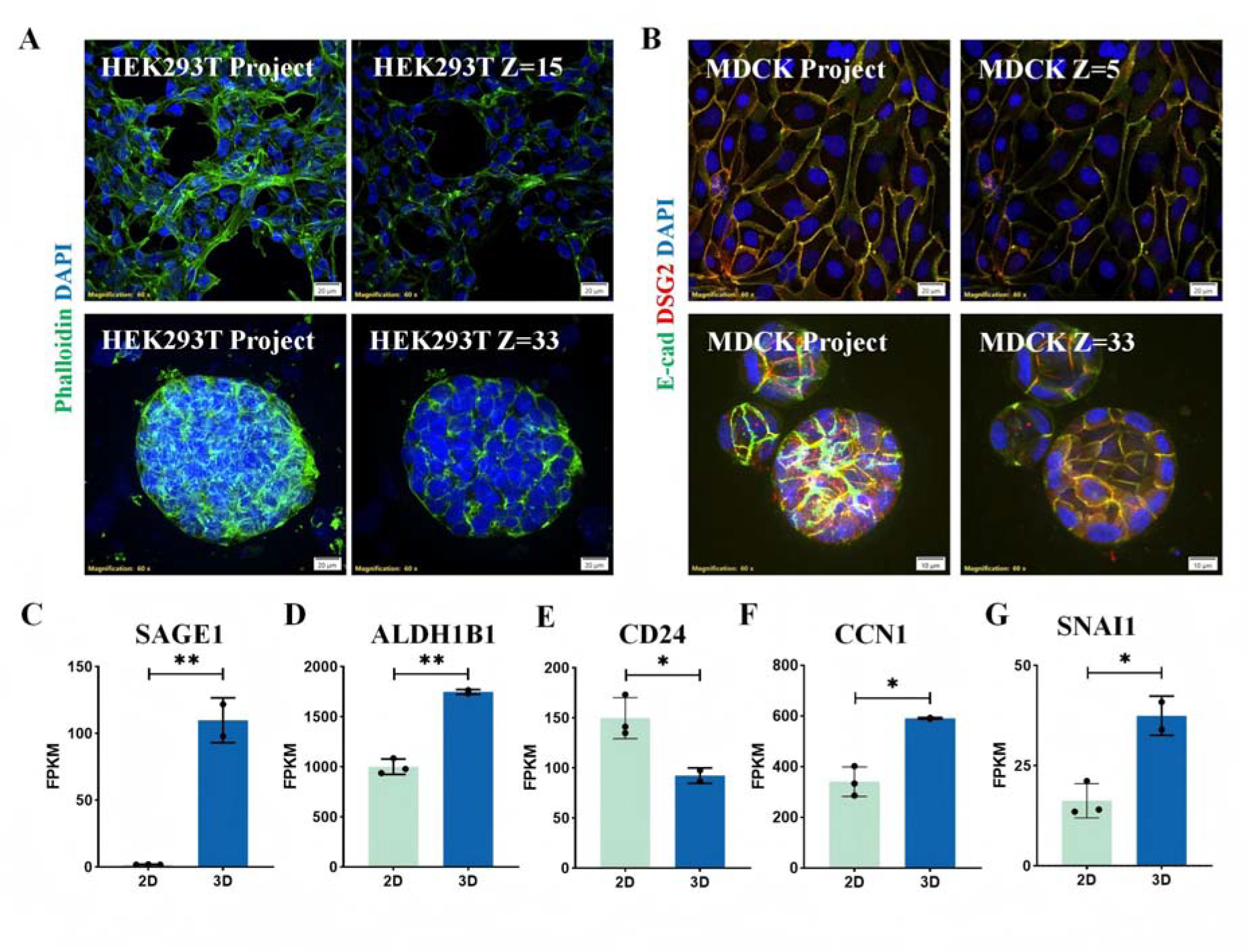
HEK293T cells form solid tumor-like spheroids and upregulate stemness/EMT markers in 3D culture. (**A**), Representative immunofluorescence confocal images of HEK293T cells cultured under 2D and 3D conditions. Upper panels: 2D culture. Left: maximum intensity projection (MIP) of all Z-stack images; right: single Z-plane image. Lower panels: 3D culture. Left: MIP of all Z-stack images; right: single Z-plane image, showing that 3D-cultured HEK293T cells form solid spheroid-like tumor structures. Cells were stained with phalloidin (F-actin, green) and DAPI (nuclei, blue). The scale bar is 20 µm. (**B**), Representative immunofluorescence confocal images of MDCK cells cultured under 2D and 3D conditions. Upper panels: 2D culture. Left: MIP of all Z-stack images; right: single Z-plane image. Lower panels: 3D culture. Left: MIP of all Z-stack images; right: single Z-plane image, showing that 3D-cultured MDCK cells form hollow tube-like structures. Cells were stained with phalloidin (F-actin, green) and DAPI (nuclei, blue). The scale bar is 20 µm. (**C-G**), Normalized gene expression levels of tumor-associated genes from RNA-seq analysis of HEK293T cells cultured under 2D vs 3D conditions: SAGE1(**C**), ALDH1B1(**D**), CD24(**E**), CCN1(**F**) and SNAI1(**G**). For all statistical plots, an unpaired Student’s t-test (n > 3 experiments, data are mean ± SD) was conducted for all data with **** P < 0.0001, *** P < 0.001, ** P < 0.1, * P <0.5, ns P > 0.5. The error bars are the standard error of the mean of the data.

Consistent with these morphological alterations, transcriptomic sequencing revealed that multiple key cancer stem cell (CSC) and epithelial–mesenchymal transition (EMT) markers were significantly upregulated in 3D-cultured HEK293T cells compared with their 2D-cultured counterparts [18, 19]. Specifically, we observed markedly increased expression of SAGE1 (a stemness-associated cancer-testis antigen), ALDH1B1 (a recognized aldehyde dehydrogenase isoform linked to CSC identity), CCN1 (a matricellular protein that promotes EMT and invasion), and SNAI1 (a core transcription factor driving EMT) (**Fig. 1C-G**). These results confirm that HEK293T cells possess a unique phenotypic plasticity and can undergo pronounced phenotypic conversion in response to different culture conditions.

In the case of 2D, both 293T and MDCK cells display an adherent monolayer phenotype, forming colonies characterized by collective cell migration. This migratory behavior differs from cancer cells, which typically migrate as individual cells. Collectively, 293T and MDCK cell lines serve as suitable models for investigating wound healing processes under 2D culture. Notably, 293T cells possess a more mesenchymal phenotype relative to MDCK cells. This observation is consistent with a growing body of literature demonstrating the remarkable lineage plasticity of HEK293 cells. Recent single-cell RNA sequencing studies have shown that HEK293 cells are most closely related to neural crest-derived adrenal medullary cells, rather than to renal epithelial cells as previously assumed [20-22]. Neural crest cells are renowned for their extraordinary migratory capacity and multipotency, properties that are essential for embryonic development [23, 24].

On this basis, we selected the standard 2D culture system—which is easy to handle and facilitates live-cell imaging and quantitative analysis—for subsequent mechanistic studies of hypoxia-induced cell migration.

### Hypoxic Stress Promotes Collective Migration and Cell Cycle Progression in HEK293T Cells

We next evaluated the functional impact of hypoxia (2% O2) on the collective migration of HEK293T cells in 2D culture using scratch wound-healing assays combined with time-lapse live-cell imaging (**Fig. 2A and C; Movie S1**). This experimental setup allowed us to continuously monitor wound closure dynamics for over 24 hours under precisely controlled oxygen conditions. HEK293T cells were labeled with LifeAct-EGFP to visualize F-actin and H2B-mCherry to mark nuclei, with imaging performed at 1 h intervals and a Z-step of 1 μm covering the entire cell height, enabling accurate capture of migratory dynamics.

**Fig. 2.**
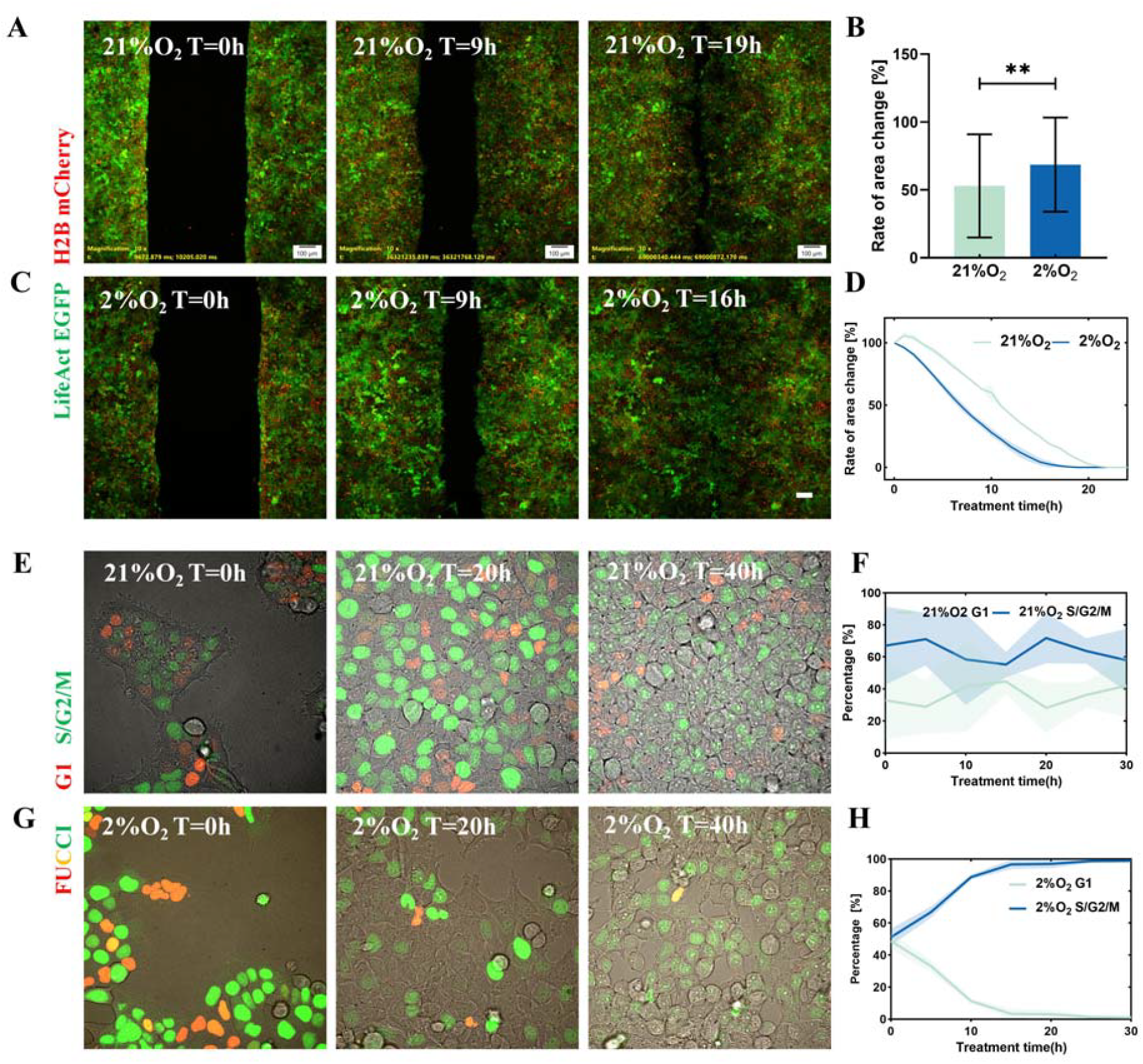
Hypoxia accelerates collective wound closure and promotes S/G2/M cell cycle progression. (**A**), Representative confocal live □cell imaging of HEK293T cells in a wound healing assay under 21% O2. Green fluorescence: F □actin labeled with LifeAct; red fluorescence: H2B □labeled nuclei. The scale bar is 100 µm. (**B**), Bar graph showing the rate of wound area change per unit time for HEK293T cells cultured under different oxygen concentrations. (**C**), Representative confocal live □cell imaging of HEK293T cells in a wound healing assay under 2% O2. Green fluorescence: LifeAct □labeled F □actin; red fluorescence: H2B. The scale bar is 100 µm. (**D**). Line graph depicting the wound area change rate over time for HEK293T cells under different oxygen concentrations. (**E**), Representative confocal live-cell imaging for cell-cycle visualization in HEK293T cells under 21% O2. Red fluorescence: hCdt1 (marking G1 phase); green fluorescence: hGem (marking S/G2/M phases). (**F**), Line graph showing the percentage of HEK293T cells in distinct cell □cycle phases over time under 21% O2. (**G**), Representative confocal live-cell imaging for cell-cycle visualization in HEK293T cells under 2% O2. Red fluorescence: hCdt1 (G1 phase); green fluorescence: hGem (S/G2/M phases). (**H**), Line graph showing the percentage of HEK293T cells in distinct cell-cycle phases over time under 2% O2. For all statistical plots, an unpaired Student’s t-test (n > 3 experiments, data are mean ± SD) was conducted for all data with **** P < 0.0001, *** P < 0.001, ** P < 0.1, * P <0.5, ns P > 0.5. The error bars are the standard error of the mean of the data.

Quantitative analysis of scratch area over time demonstrated that hypoxic exposure significantly accelerated wound closure, with a substantially higher wound-healing rate compared with the normoxic control group (21% O2) (**Fig. 2B and D**). At 16 h post-scratch, cultures in the hypoxic group had essentially completed wound closure, whereas the normoxic group achieved closure only by 20 h.

The increased wound closure rate under hypoxia could stem from two sources: enhanced single-cell migration speed and/or increased cell numbers contributing to faster healing [25, 26]. To determine whether proliferation contributed to the accelerated scratch closure under hypoxia, we employed the FUCCI (fluorescence ubiquitination-based cell cycle indicator) system [27], which enables real-time observation of cell cycle phases in living cells (**Fig. 2E and G; Movie S1**). Interestingly, tracking of cell cycle dynamics revealed a significantly elevated proportion of cells in S/G2/M phases under hypoxic conditions, indicating that local hypoxic stress paradoxically accelerated the cell division cycle, thereby jointly propelling the advancement of the wound-healing front (**Fig. 2F and H**). After 40 h of hypoxia, nearly 100% of cells were in S/G2/M phases, whereas under normoxia the proportion was approximately 60% at the same time point (**Fig. 2F and H**). These results indicate that hypoxia enhances the wound-healing capacity of HEK293T cells in 2D culture by concurrently increasing cell proliferation.

We propose that this mesenchymal phenotype and lineage plasticity (e.g., stemness) [17] of HEK293T cells serves as the intrinsic basis for their robust migratory response under hypoxic conditions. Unlike terminally differentiated epithelial cells, HEK293T cells retain the ability to reactivate embryonic developmental programs, enabling rapid phenotypic switching and acquisition of enhanced migratory capacity in response to microenvironmental stresses such as hypoxia [28-30]. The solid tumor-like spheroid morphology and upregulated stemness gene expression observed in 3D culture exemplify this plasticity, while the hypoxia-induced migration we observed in the 2D culture system represents a manifestation of the same plasticity under a different microenvironmental stress [18-20].

### Hypoxia Augments Single-Cell Motility and Induces Morphological Spreading

To resolve the single-cell behavioral changes underlying collective migration, we performed morphometric analysis of bright-field images (**Fig. 3A**). Under hypoxic conditions, HEK293T cells in 2D culture displayed a more spread and elongated morphology, characterized by significantly increased surface area and elevated eccentricity, suggesting lamellipodial extension and cytoskeletal remodeling (**Fig. 3B-C**). Quantitative analysis showed that the average cell area of hypoxic cells was more than 30% larger than that of normoxic cells, and eccentricity was 10% higher (**Fig. 3B-C**) - morphological changes closely associated with enhanced migratory capacity.

**Fig. 3.**
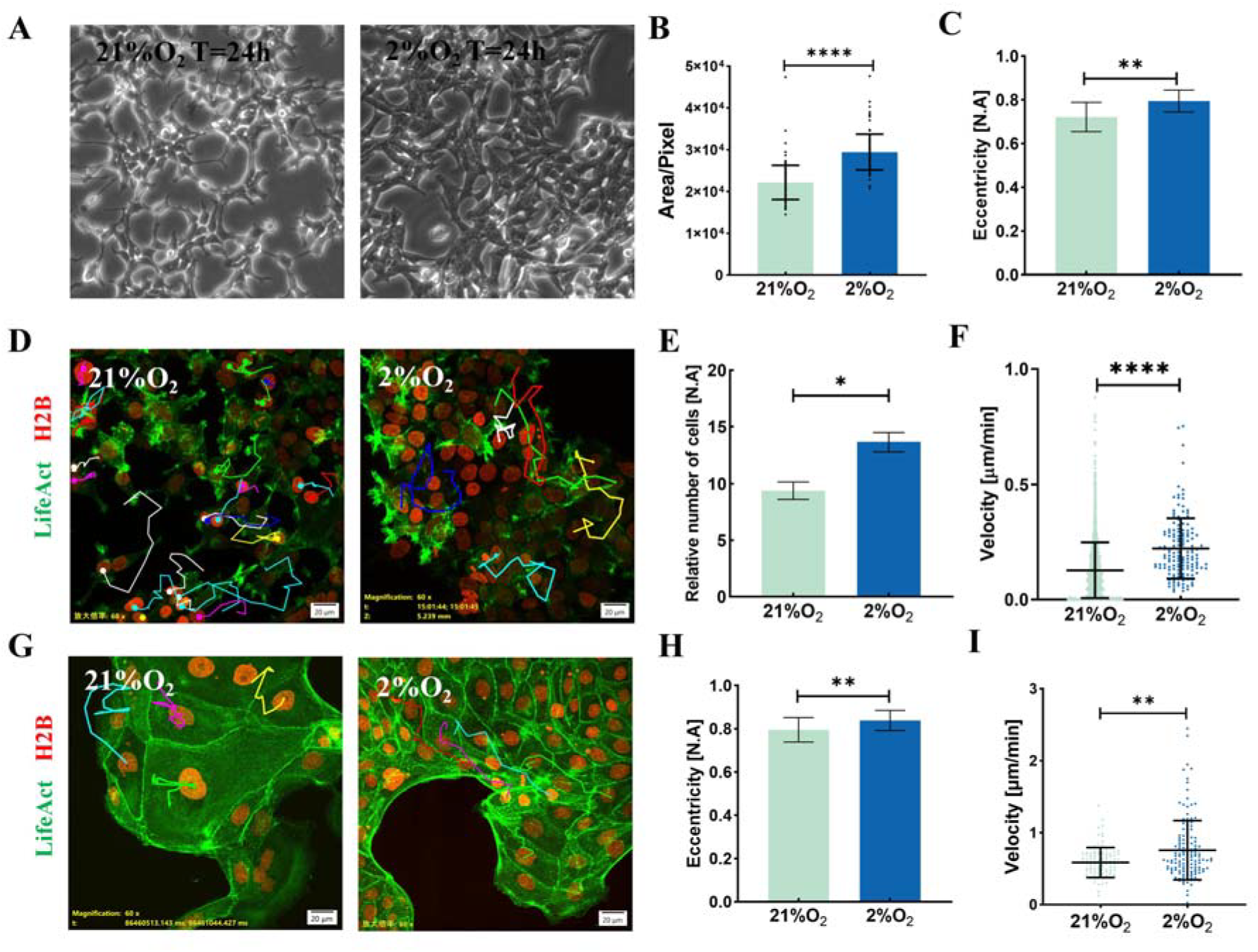
Hypoxia induces morphological spreading and enhances single-cell migration velocity. (**A**), Representative bright-field images of HEK293T cells cultured under normoxia (left) and hypoxia (right) for 24 h. (**B**), Quantification of cell area under different culture conditions. (**C**), Quantification of HEK293T cell eccentricity under different culture conditions. (**D**), Representative confocal images of HEK293T cell migration tracks under normoxia (left) and hypoxia (right). F-actin was visualized with LifeAct-EGFP (green), and nuclei were stained with H2B-mCherry (red). The scale bar is 20 µm. (**E**), Quantification of relative cell number under different culture conditions. (**F**), Quantification of HEK293T cell migration velocity under different culture conditions. (**G**), Representative confocal images of MDCK cell migration tracks under normoxia (left) and hypoxia (right). F-actin was visualized with LifeAct-EGFP (green), and nuclei were stained with H2B-mCherry (red). The scale bar is 20 µm. (**H**), Quantification of MDCK cell eccentricity under different culture conditions. (**I**), Quantification of MDCK cell migration velocity under different culture conditions. For all statistical plots, an unpaired Student’s t-test (n > 3 experiments, data are mean ± SD) was conducted for all data with **** P < 0.0001, *** P < 0.001, ** P < 0.1, * P <0.5, ns P > 0.5. The error bars are the standard error of the mean of the data.

High-resolution tracking using 60× oil-immersion time-lapse microscopy corroborated these findings, capturing a marked increase in both the migration speed and cell density of individual cells at the migratory front (**Fig. 3D-E**). Single-cell tracking analysis revealed that the mean instantaneous velocity of hypoxic cells was approximately 1.7-fold that of normoxic cells (**Fig. 3F**). These results demonstrate that hypoxia promotes wound healing by enhancing the migratory ability of individual cells. MDCK cells are a well-established model for investigating the mechanisms of wound healing [11], see the purse string in Fig.3G. To further explore how hypoxia promotes cell motility and spreading, thereby regulating wound healing, we repeated the above experiments using MDCK cells (**Fig. 3G-I**). Morphological analysis revealed that hypoxia increased the eccentricity of MDCK cell morphology (**Fig. 3H**). Furthermore, single-cell trajectory analysis demonstrated that hypoxia significantly enhanced cell velocity by approximately 20% (**Fig. 3I**).

Under hypoxic microenvironments, the average instantaneous velocity of single HEK293T and MDCK cells is significantly elevated, accompanied by pronounced morphological spreading and elongation. This morphological remodeling is not merely a superficial phenotypic shift, but rather a geometric manifestation of highly optimized cell migration mechanics [31]. From a biomechanical perspective, the spreading of cell morphology substantially expands the effective contact area with the extracellular matrix (ECM). This expansion provides a broader geometric space for the assembly of focal adhesions and the mechanotransduction of integrins, such as ITGB3, thereby enhancing traction forces at the leading edge [32, 33]. Concurrently, the increase in eccentricity marks the successful establishment of morphological cell polarity. This symmetry-breaking event prompts internal stress fibers to align parallel to the long axis of the cell, converting the contractile forces generated by myosin from an isotropic state into a directionally focused orientation, which maximizes the net resultant force along the direction of migration [34, 35]. This process fundamentally relies on protrusion extension and intensive remodeling of the actin cytoskeleton at the microscopic scale [36]. The perfect synergy between morphology and mechanics collectively constitutes the physical and biological foundation by which hypoxia accelerates epithelial wound healing.

### Down-regulation of Desmoglein-2 (DSG2) Weakens Intercellular Cohesion Under Hypoxia

Given that cell migration requires the precise coordination of cell-cell detachment [37, 38], we used live-cell fluorescence imaging to dynamically track the desmosomal cadherin desmoglein-2 (DSG2) (**Fig. 4A and C; Movie S3**). We labeled HEK293T cells with DSG2-mCherry. DSG2 is a core component of desmosomes, specialized intercellular junctions that provide strong mechanical adhesion between epithelial cells. We observed that the fluorescence signal intensity of DSG2 in 2D-cultured HEK293T cells was persistently downregulated during hypoxic exposure (**Fig. 4B and D**). Using the same method, we visualized DSG2 expression in MDCK cells. The results showed that under normoxic conditions, the fluorescence signal of DSG2 remained relatively stable (**Fig. 4E and F**), whereas under hypoxic conditions, it decreased in a time-dependent manner (**Fig. 4G and H**).

**Fig. 4.**
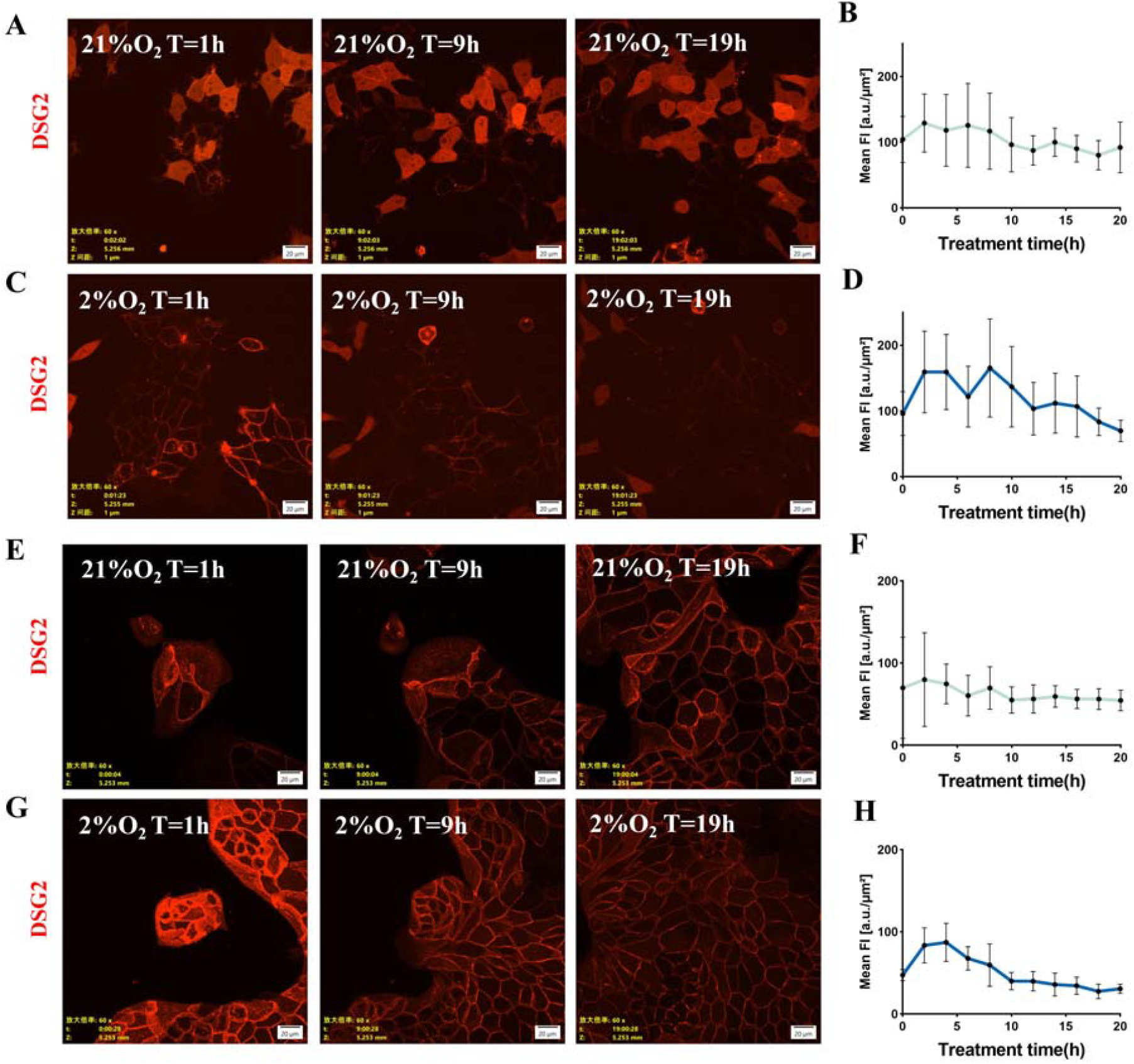
Hypoxia persistently downregulates DSG2 expression and weakens intercellular cohesion. (**A**), Representative confocal images from live-cell imaging of DSG2 expression in HEK293T cells at the indicated time points under normoxic conditions. The scale bar is 20 µm. (**B**), Quantification of DSG2 fluorescence intensity over time under normoxia. (**C**), Representative confocal images from live-cell imaging of DSG2 expression in HEK293T cells at the indicated time points under hypoxic conditions. The scale bar is 20 µm. (**D**), Quantification of DSG2 fluorescence intensity over time under hypoxia. (**E**), Representative confocal images from live-cell imaging of DSG2 expression in MDCK cells at the indicated time points under normoxic conditions. The scale bar is 20 µm. (**F**), Quantification of DSG2 fluorescence intensity over time under normoxia. (**G**), Representative confocal images from live-cell imaging of DSG2 expression in MDCK cells at the indicated time points under hypoxic conditions. The scale bar is 20 µm. (**H**), Quantification of DSG2 fluorescence intensity over time under hypoxia. For all statistical plots, an unpaired Student’s t-test (n > 3 experiments, data are mean ± SD) was conducted for all data with **** P < 0.0001, *** P < 0.001, ** P < 0.1, * P <0.5, ns P > 0.5. The error bars are the standard error of the mean of the data.

This rapid disassembly of DSG2-mediated intercellular junctions is directly consistent with the observed morphological spreading, suggesting that hypoxia promotes the dissociation of stable epithelial-like sheets to facilitate flexible cell movement. From a mechanobiological perspective, desmosomes exert a powerful lateral constraint that preserves collective confinement and stabilizes cortical tension at cell-cell boundaries [39]. The persistent downregulation of DSG2 under hypoxia essentially unloads this intercellular tension, lowering the physical energy barrier required for individual cells to escape the cohesive sheet. This reduction in intercellular tethering likely liberates intracellular cytoskeletal resources, allowing G-actin to reallocate toward the basal plane to drive focal adhesion assembly and lamellipodial protrusion. The downregulation of DSG2 expression is expected to weaken cell-cell adhesion, enabling individual cells to more readily detach from the epithelial sheet and migrate into the wound area [40, 41].

DSG2 is a core component of desmosomes and is responsible for maintaining the structural integrity of epithelial sheets [39]. In a normal epithelial sheet, adjacent cells are tightly locked by desmosomes, establishing a state of high collective confinement. Within this constrained microenvironment, even if individual cells possess an intrinsic propensity for locomotion, their surrounding physical barriers severely restrict their degrees of freedom [42]. The persistent downregulation of DSG2 does not merely disrupt localized intercellular junctions; rather, it physically dismantles this “spatial confinement” and reduces the shear resistance of the epithelial sheet [43]. Consequently, this structural dissolution lowers the activation energy barrier for single-cell escape, directly clearing the spatial obstacles required for subsequent acceleration of single-cell velocity and morphological spreading.

The sustained downregulation of DSG2 under hypoxia weakens cell–cell adhesion, allowing cells to detach from their neighbors and migrate individually or in small groups. This finding is consistent with previous studies in breast cancer, where loss of DSG2 has been shown to promote EMT and enhance the invasive capacity of cancer cells, suggesting that this may be a general mechanism by which hypoxia facilitates cell migration [40].

### Biphasic Dynamics of Integrin β3(ITGB3) Under Hypoxia

Concurrently, live-cell tracking of the cell-matrix adhesion molecule integrin β3 (ITGB3) in HEK293T cells revealed a unique biphasic kinetic response that emerged specifically under hypoxic stress (**Fig. 5A-C; Movie S4**). During the initial hours of hypoxia (0-3 h), the ITGB3 fluorescence intensity in 2D-cultured HEK293T cells rose sharply above normoxic levels, presumably to reinforce focal adhesions required for initial traction force generation (**Fig. 5D**). Subsequently, the ITGB3 signal underwent a robust time-dependent downregulation, falling significantly below the normoxic baseline by 9 h and continuing to decline through 24 h (**Fig. 5D**). This biphasic pattern indicates that cell-matrix adhesion is subjected to precise temporal control to optimize cell migration efficiency.Transcriptome sequencing was performed on HEK293T cells cultured under normoxic or hypoxic conditions for 24 hours. The normalized expression levels of ITGB1(**Fig. 5E**), ITGB2(**Fig. 5F**), ITGB3(**Fig. 5G**), and ITGB3BP (**Fig. 5H**) were compared between the two culture conditions. The results revealed a downward trend in the expression levels of these genes under hypoxic conditions.

**Fig. 5.**
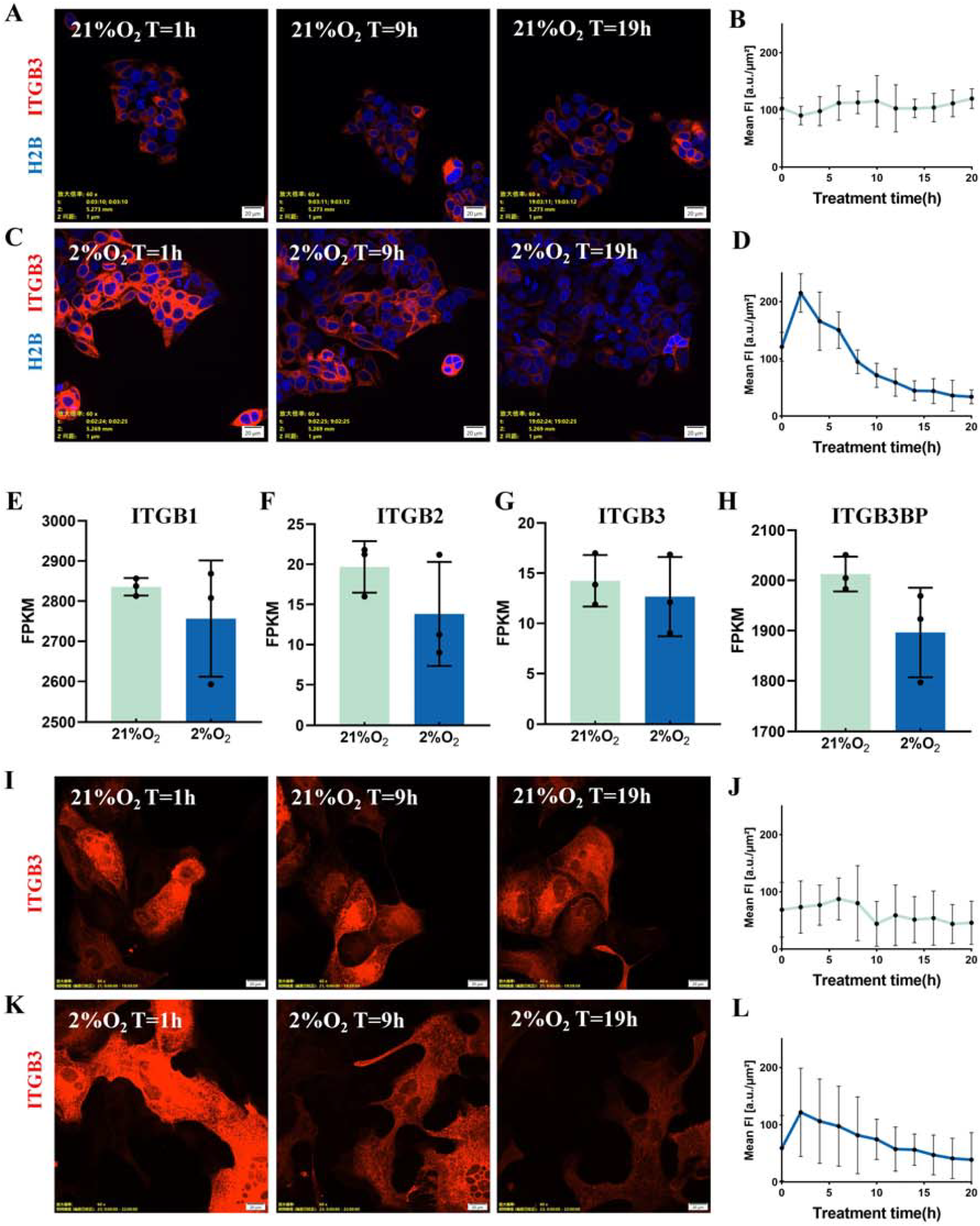
ITGB3 exhibits a biphasic expression pattern under hypoxic stress. (**A**), Representative confocal images from live-cell imaging of ITGB3 expression in HEK293T cells at the indicated time points under normoxic conditions. The scale bar is 20 µm. (**B**), Quantification of ITGB3 fluorescence intensity over time under normoxia. (**C**), Representative confocal images from live-cell imaging of ITGB3 expression in HEK293T cells at the indicated time points under hypoxic conditions. The scale bar is 20 µm. (**D**), Quantification of ITGB3 fluorescence intensity over time under hypoxia. (**E–H**) Normalized expression levels of cell–extracellular matrix interaction-related genes, including ITGB1 (**E**), ITGB2 (**F**), ITGB3 (**G**), and ITGBBP (**H**), obtained from transcriptomic sequencing of HEK293T cells cultured under different conditions. (**I**), Representative confocal images from live-cell imaging of ITGB3 expression in MDCK cells at the indicated time points under normoxic conditions. The scale bar is 20 µm. (**J**), Quantification of ITGB3 fluorescence intensity over time under normoxia. (**K**), Representative confocal images from live-cell imaging of ITGB3 expression in MDCK cells at the indicated time points under hypoxic conditions. The scale bar is 20 µm. (**L**), Quantification of ITGB3 fluorescence intensity over time under hypoxia. For all statistical plots, an unpaired Student’s t-test (n > 3 experiments, data are mean ± SD) was conducted for all data with **** P < 0.0001, *** P < 0.001, ** P < 0.1, * P <0.5, ns P > 0.5. The error bars are the standard error of the mean of the data.

Long-term live-cell imaging of ITGB3 in MDCK cells yielded results consistent with those observed in HEK293T cells (**Fig. 5I-L**). Under normoxic conditions, the fluorescence signal of ITGB3 in MDCK cells remained relatively stable (**Fig. 5I and J**). In contrast, under hypoxic conditions, ITGB3 fluorescence expression was significantly upregulated during the first few hours of hypoxia, but subsequently declined with prolonged hypoxic exposure, eventually falling below the signal intensity observed under normoxic conditions (**Fig. 5K and L**).

The morphological spreading of HEK293T cells under hypoxia correlates with the initial upregulation of ITGB3. Specifically, the rapid increase of ITGB3 during the early phase of hypoxia provides a substantial number of focal adhesions that stabilize the spread morphology and generate robust initial traction forces. As ITGB3 levels subsequently decline in the late phase, coupled with the persistent downregulation of DSG2, cells avoid being immobilized by excessive cell-matrix adhesion. Conversely, this moderate reduction in cell-matrix adhesion facilitates the successful detachment of the trailing edge. This coordinated remodeling allows cells to maintain a streamlined, high-eccentricity morphology, thereby achieving efficient “slippage-based” migration.

In contrast to the monotonic downregulation of DSG2, ITGB3 exhibited a striking biphasic expression pattern under hypoxia. The upregulation of ITGB3 during the initial 3 h of hypoxia likely serves to reinforce focal adhesions and provide the traction force necessary for cells to pull themselves forward. This is consistent with prior studies showing that hypoxia upregulates ITGB3 expression in a HIF-1α-dependent manner and that ITGB3 is required for hypoxia-induced migration in multiple cell types [44, 45]. However, the subsequent downregulation of ITGB3 after 3 h is unexpected and has not been widely reported. We propose that this late-phase downregulation is critical for efficient cell migration, as it allows cells to detach their trailing edge from the matrix and move forward. Persistently high levels of ITGB3 would lead to excessive adhesion, thereby impeding cell migration. Thus, this biphasic regulation of ITGB3 represents a sophisticated mechanism for optimizing adhesion dynamics during migration.

### Transcriptomic Profiling Validates Hypoxia-Induced Phenotypic and Adhesion Remodeling

To obtain a global overview of the molecular landscape governing the observed phenotypic adaptations, we performed RNA sequencing (RNA-seq) on 2D-cultured HEK293T cells under both normoxic and hypoxic conditions. Enrichment analysis of differentially expressed genes (DEGs), comprising Gene Ontology (GO) terms and Kyoto Encyclopedia of Genes and Genomes (KEGG) pathways, robustly coincided with our in vitro experimental observations.

Specifically, Biological Process (BP) analysis revealed significant enrichment in terms related to “cell–cell adhesion via plasma-membrane adhesion molecules” and “extracellular matrix/structure organization” (**Fig. 6A**). This transcriptomic profile directly supports our experimental findings that hypoxia dynamically perturbs cell-cell cohesion—evidenced by the continuous downregulation of the desmosomal cadherin DSG2—and induces prominent cell flattening and morphological extension. Furthermore, the prominent enrichment of processes governing intracellular ions, such as “regulation of cytosolic calcium ion concentration” and “adenylate cyclase-modulating G protein-coupled receptor signaling pathway”, echoes the accelerated cell cycle progression and heightened metabolic activity observed under hypoxic stress.

**Fig. 6.**
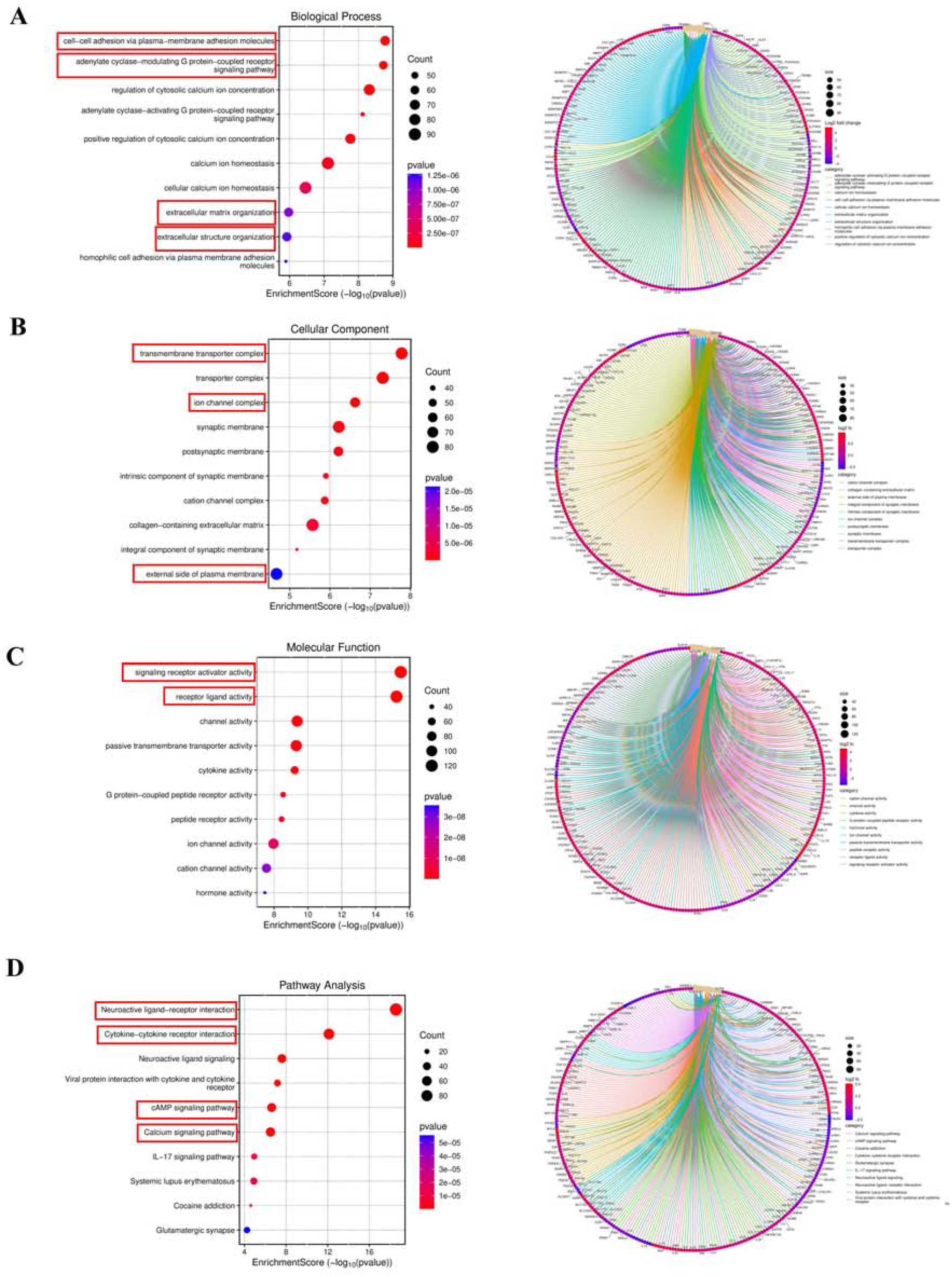
Transcriptome sequencing enrichment analysis. RNA-seq was performed on HEK293T cells cultured under normoxia or hypoxia for 24 h. Differentially expressed genes (DEGs) were identified and subjected to Gene Ontology (GO) and KEGG pathway enrichment analyses. (**A**), Dot plot of enrichment scores for Biological Process (BP) terms (Left). Dot size represents gene count, and color intensity indicates the adjusted P value. Cnetplot depicting DEGs associated with enriched Biological Process (BP) terms (Right). Node colors denote log □ fold changes. (**B**), Dot plot of enrichment scores for Cellular Component (CC) terms (Left). Cnetplot depicting DEGs associated with enriched Cellular Component (CC) terms (Right). (**C**), Dot plot of enrichment scores for Molecular Function (MF) terms (Left). Cnetplot depicting DEGs associated with enriched Molecular Function (MF) terms (Right). (**D**), Dot plot of enrichment scores for KEGG pathway terms (Left). Cnetplot depicting DEGs associated with enriched KEGG pathway terms (Right).

In alignment with these biological processes, Cellular Component (CC) analysis localized the responsive gene products predominantly to the “transmembrane transporter complex”, “ion channel complex”, and the “external side of plasma membrane” (**Fig. 6B**). This spatial distribution strongly correlates with the profound reorganization of adhesion and cytoskeletal machinery at the cell boundaries, providing a structural basis for the biphasic expression pattern of the cell-matrix adhesion molecule ITGB3 and the dismantling of DSG2 junctions.

Consistently, Molecular Function (MF) analysis identified “signaling receptor activator activity” and “receptor ligand activity” as the most prominent nodes, accompanied by substantial enrichment in “channel activity” and “passive transmembrane transporter activity” (**Fig. 6C**). This functional repertoire suggests a heightened sensitivity of hypoxic cells to microenvironmental cues.

Finally, Pathway Analysis (PA) via KEGG mapping further integrated these molecular signatures into highly convergent networks, primarily driven by “Neuroactive ligand–receptor interaction”, “Cytokine–cytokine receptor interaction”, and the “cAMP/Calcium signaling pathways” (**Fig. 6D**). Rather than pointing to a singular linear cascade, these co-enriched pathways collectively reflect a state of intensive environmental sensing and ion homeostasis adjustments. Taken together, these transcriptomic enrichment data independently corroborate our cellular and morphological findings, demonstrating that hypoxia triggers a coordinated transcriptional program aimed at downregulating stable cell-cell junctions and modulating cell-matrix interactions to facilitate phenotypic adaptation.

## Discussion

In this study, we demonstrated that HEK293T cells possess a unique phenotypic plasticity, exhibiting solid tumor-like spheroid morphology and stemness gene expression signatures in 3D culture that are clearly distinct from classical epithelial cells. But in 2D culture, 293T and MDCK cells exhibit adherent monolayers with collective migration, distinct from individually migrating cancer cells. Both lines are appropriate wound healing models, yet 293T cells show stronger mesenchymal traits than MDCK cells.

Building on this, we systematically investigated the impact of hypoxia on HEK293T and MDCK cell migration in a standard 2D culture system and found that hypoxia significantly accelerated both collective and single-cell migration, accompanied by cell cycle acceleration and morphological spreading. Mechanistically, we discovered that hypoxia dynamically regulates two key adhesion molecules: DSG2 undergoes persistent downregulation to weaken cell-cell adhesion, while ITGB3 displays a distinctive biphasic expression pattern to optimize cell-matrix adhesion required for migration. Transcriptomic profiling further corroborated these phenotypic transitions, revealing a global enrichment of gene networks governing plasma-membrane adhesion, receptor-ligand activities, and ion homeostasis that independently mirrors the observed junctional remodeling and accelerated cellular responses. These findings provide new insights into the molecular mechanisms linking oxygen sensing to cell adhesion and migration, with important implications for both wound healing and cancer metastasis.

Our most important mechanistic insight is the discovery that hypoxia promotes cell migration by coordinately regulating cell-cell and cell-matrix adhesion. This mechanism relies on the precise temporal control of DSG2 and ITGB3 expression to balance the two opposing forces of intercellular cohesion and cell-matrix traction. The cooperative nature of these two processes-DSG2 downregulation weakening cell-cell adhesion and biphasic ITGB3 kinetics optimizing cell-matrix adhesion-enables cells to move efficiently through the extracellular matrix. This “adhesion-slippage” mechanism has been proposed to explain the high migratory capacity of certain cancer cells [40, 46], and our live-cell imaging data provide direct visual evidence for this model in the context of hypoxia-induced migration.

## Summary

In conclusion, our study reveals a novel mechanism by which hypoxia accelerates wound-healing migration in 2D-cultured HEK293T and MDCK cells. This mechanism involves the cooperative downregulation of DSG2 to weaken intercellular adhesion and the biphasic regulation of ITGB3 to optimize cell-matrix adhesion, a coordinated process that is comprehensively supported by transcriptomic profiling showing global alterations in membrane adhesion networks and intracellular ion homeostasis. These findings not only advance our understanding of the molecular mechanisms underlying hypoxia-induced cell migration but also provide potential therapeutic targets for promoting wound healing and inhibiting cancer metastasis. Future studies are needed to validate these findings in vivo and to further dissect the precise biophysical cues and upstream microenvironmental factors that initiate this dramatic remodeling of the DSG2 and ITGB3 adhesion molecules.

## Acknowledgments

We thank Ruirui Gao and Yeqing Leng from Wuhan University for sharing monoclonal transfected cell lines and DeepSeek V4 for grammar checking and text polishing.

## Funding

National Natural Science Foundation of China (NSFC): Bo Li Distinguished Young Scholars Grant (Overseas)

Wuhan University (WHU): Bo Li Talents Startup Funding

National Natural Science Foundation of China (NSFC): Bo Li Young Scientists Fund (C Class)

## Author contributions

Conceptualization: BL

Methodology: BL, ZDW

Investigation: BL, ZDW, LT

Visualization: ZDW

Funding acquisition: BL

Project administration: BL

Supervision: BL

Writing – original draft: ZDW, BL

Writing – review & editing: ZDW, BL, LT

## Competing interests

The Authors declare that they have no competing interests.

## Data and materials availability

All data are available in the main text or the supplementary materials.

## Supplementary Materials

Legends for Movies S1 to S4

## Legends for Supplementary Movies

**Movie S1. Wide-field time-lapse confocal imaging was performed to monitor wound healing in HEK293T cells cultured under 21% O2(left) and 2% O2(right) condition**. A 10× objective lens was ued. F-actin was labeled with Lifeact-EGFP (excitation at 488 nm; laser power 20%; exposure time 200 ms), and nuclei were labeled with H2B-mCherry (excitation at 561 nm; laser power 20%; exposure time 200 ms). Images were acquired every 1 hour for a total duration of 28 hours. Scale bar = 100 μm.

**Movie S2. Wide-field time-lapse confocal imaging was performed to visualize the cell cycle in EK293T cells cultured under 21% O2(left) and 2% O2(right) condition using the FUCCI system**. A 60× oil immersion objective lens was used. Cells in S/G2/M phases were labeled with green fluorescent potein fused to Geminin (excitation at 488 nm; laser power 20%; exposure time 200 ms), and cells in G1 phase were labeled with red fluorescent protein fused to Cdt1 (excitation at 561 nm; laser power 20%; exposure time 200 ms). Images were acquired every 1 hour for a total duration exceeding 40 hours. Scale bar = 20μm.

**Movie S3. Wide-field time-lapse confocal imaging was performed to visualize DSG2 expression in HEK293T cells cultured under 21% O2(left) and 2% O2(right) condition**. A 60× oil immersion objective lens was used. DSG2 was labeled with DSG2-mCherry (excitation at 561 nm; laser power 20%; exposure time 200 ms). Images were acquired every 1 hour for a total duration of 24 hours. Scale bar = 20 μm.

**Movie S4. Wide-field time-lapse confocal imaging was performed to visualize ITGB3 expression in HEK293T cells cultured under 21% O2(left) and 2% O2(right) condition**. A 60× oil immersion objective lens was used. ITGB3 was labeled with ITGB3-mCherry (excitation at 561 nm; laser power 20%; exposure time 200 ms). Images were acquired every 1 hour for a total duration of 24 hours. Scale bar = 20 μm.

## Notes

### Competing Interest Statement

The authors have declared no competing interest.

