## Supplementary figures and images for "Hypoxia Promotes Wound Healing via Dynamical-Mechanical Balance and Adhesion Remodeling"

### Figure 1

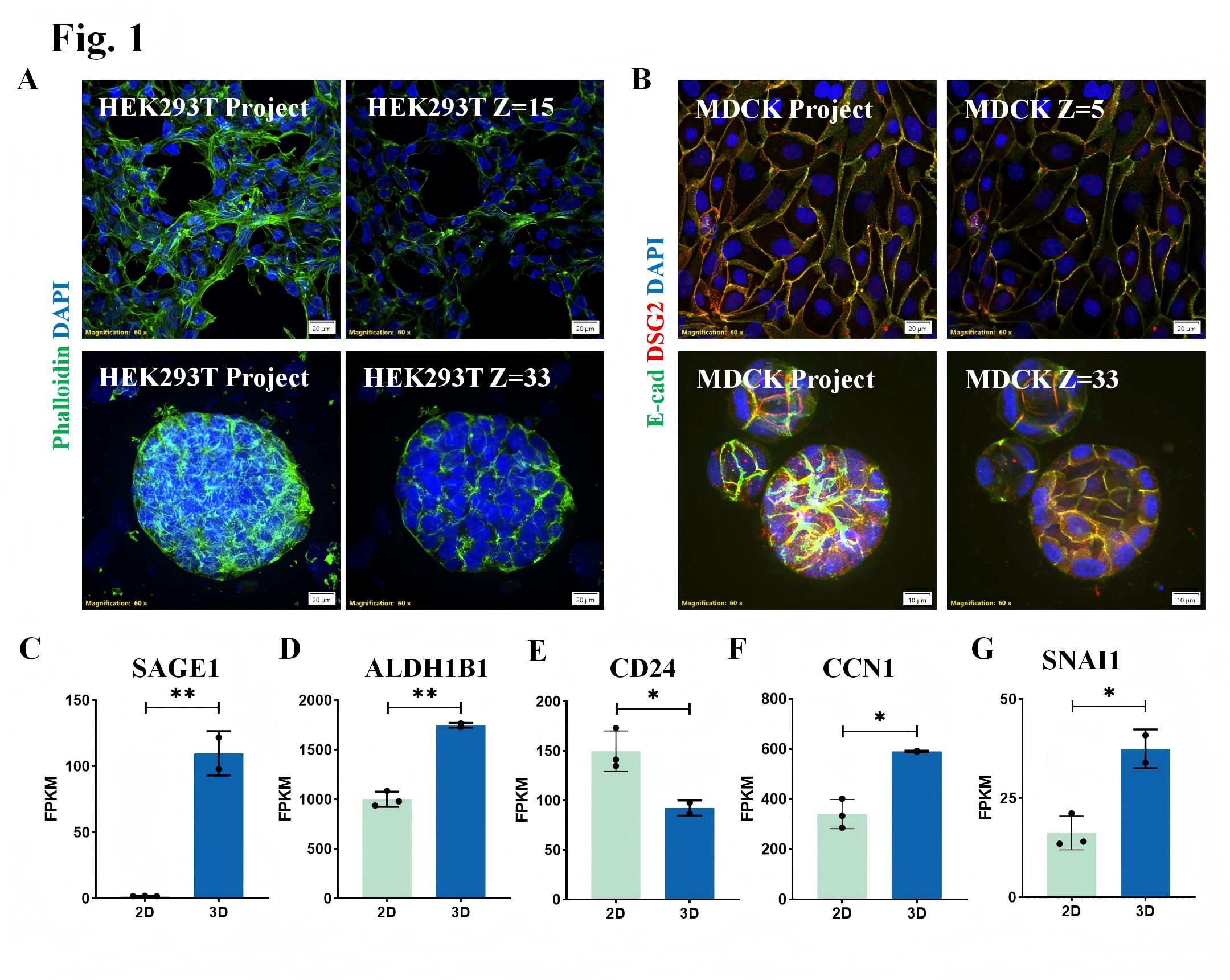

### Figure 2

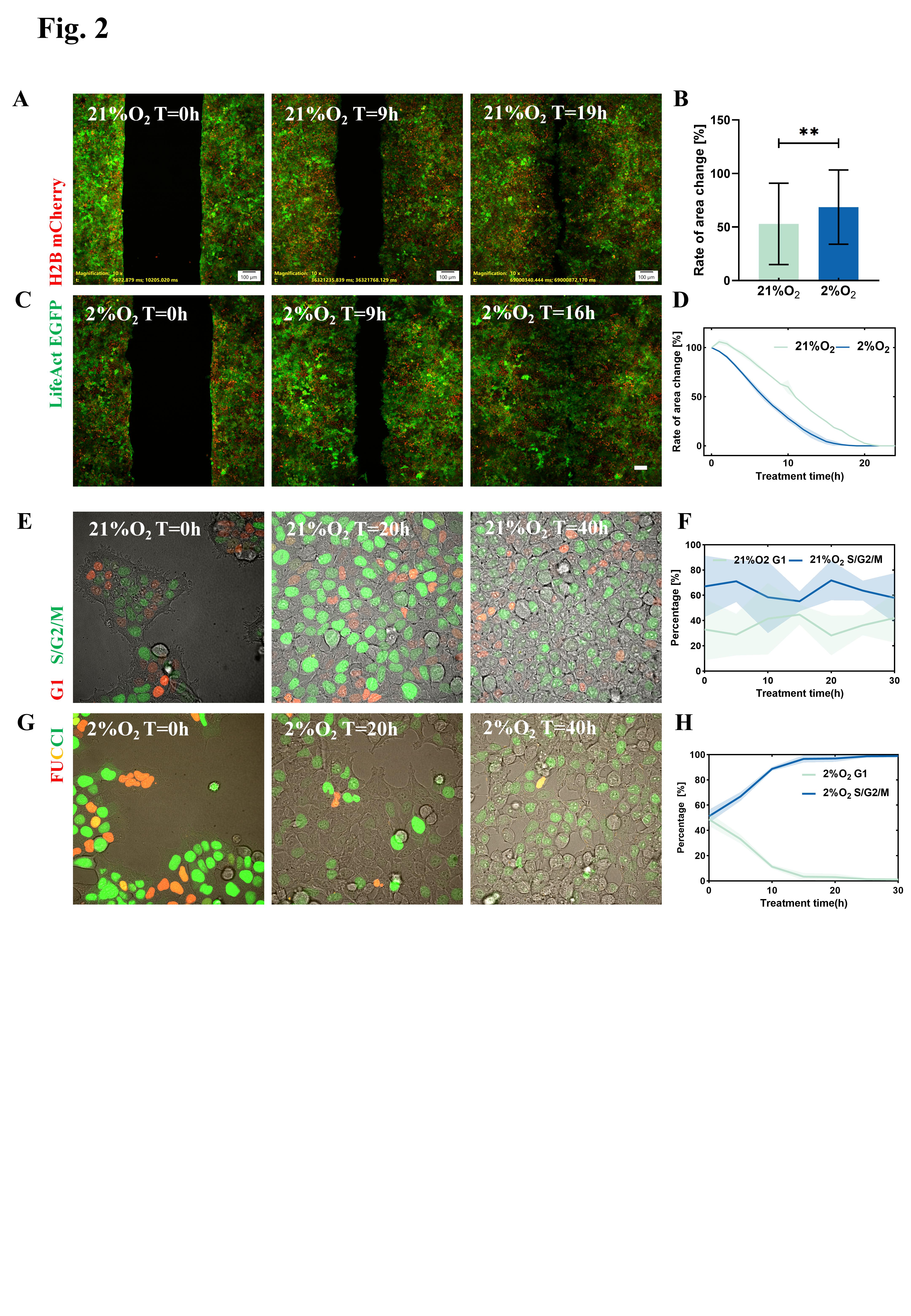

### Figure 3

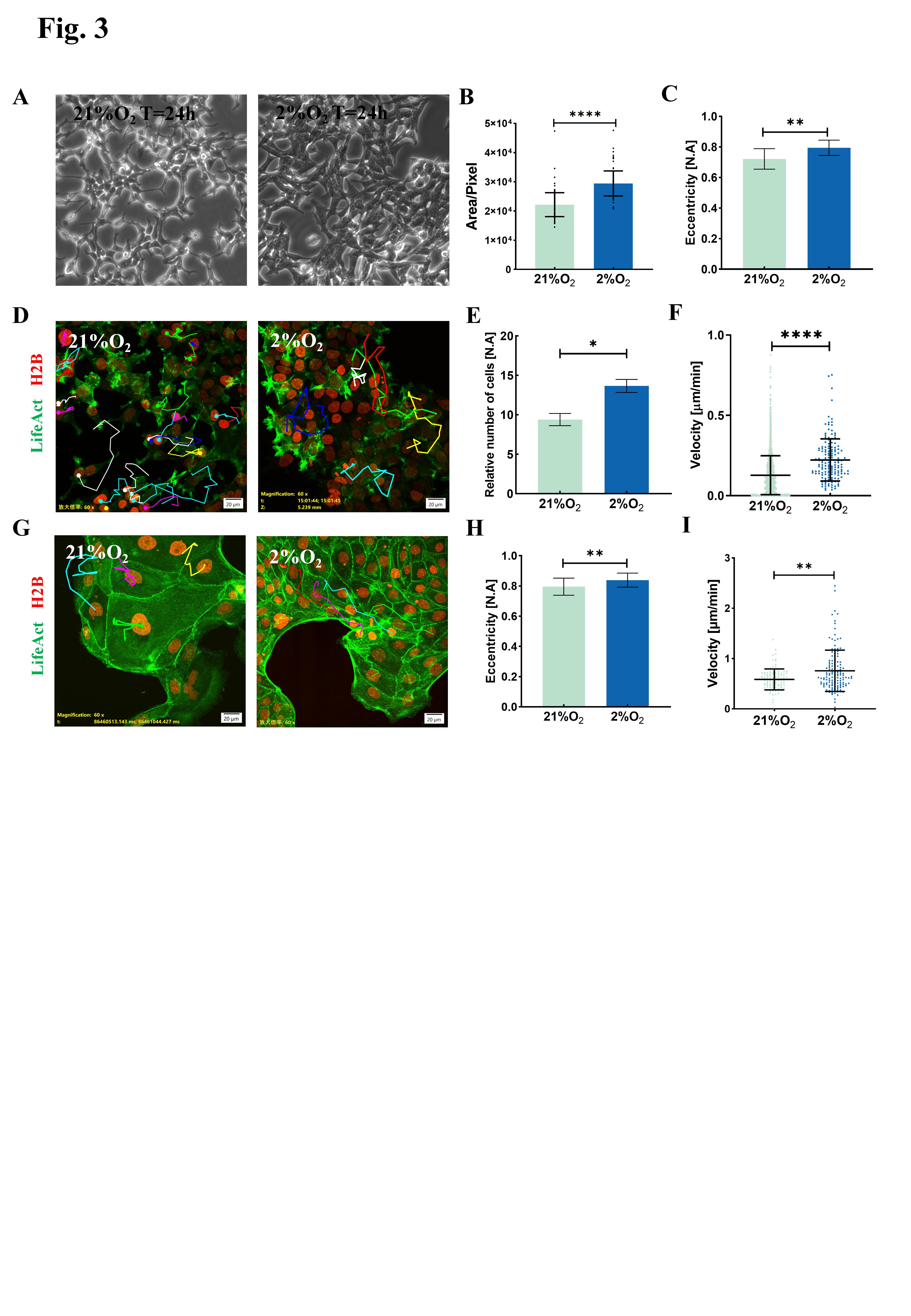

### Figure 4

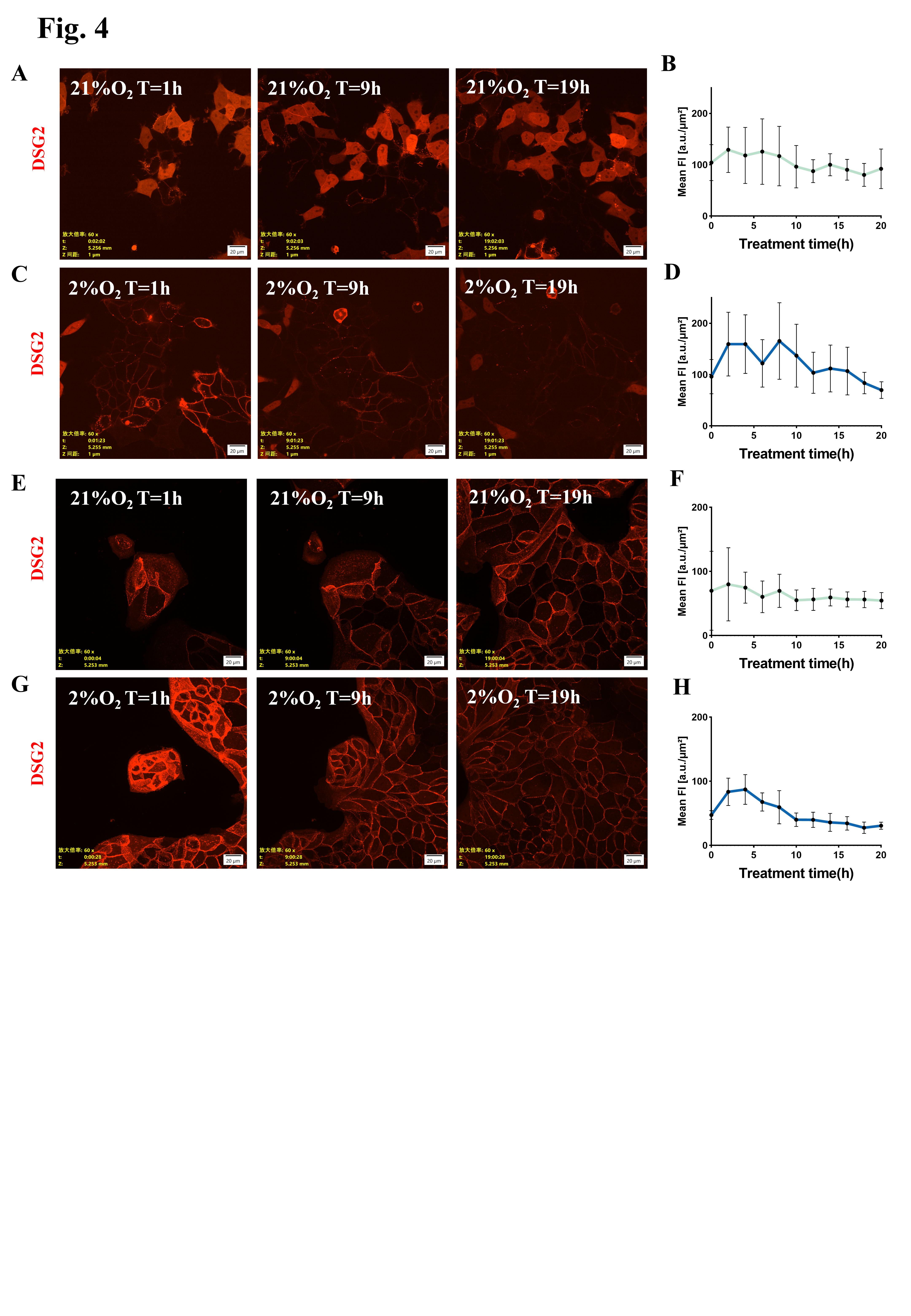

### Figure 5

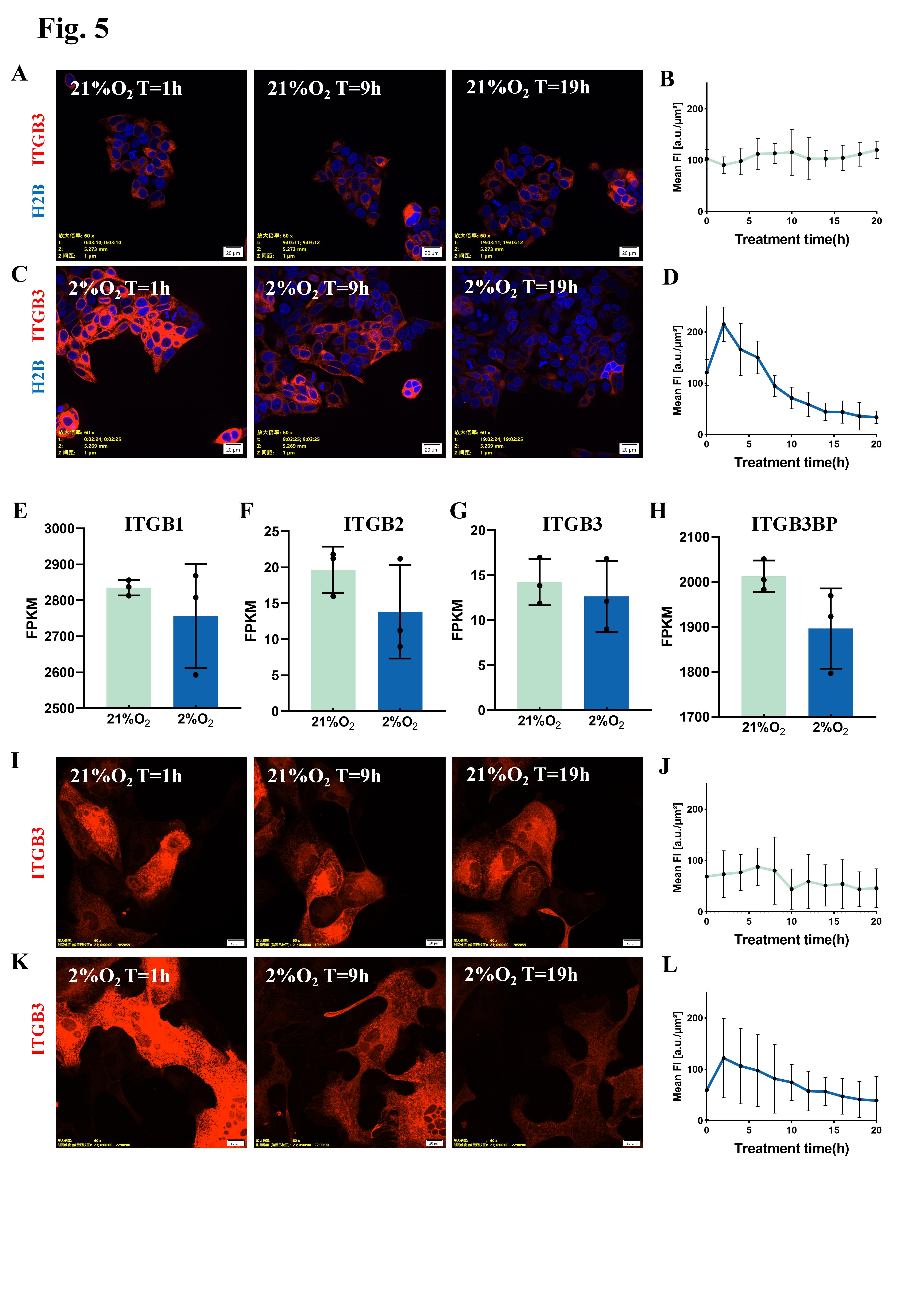

### Figure 6

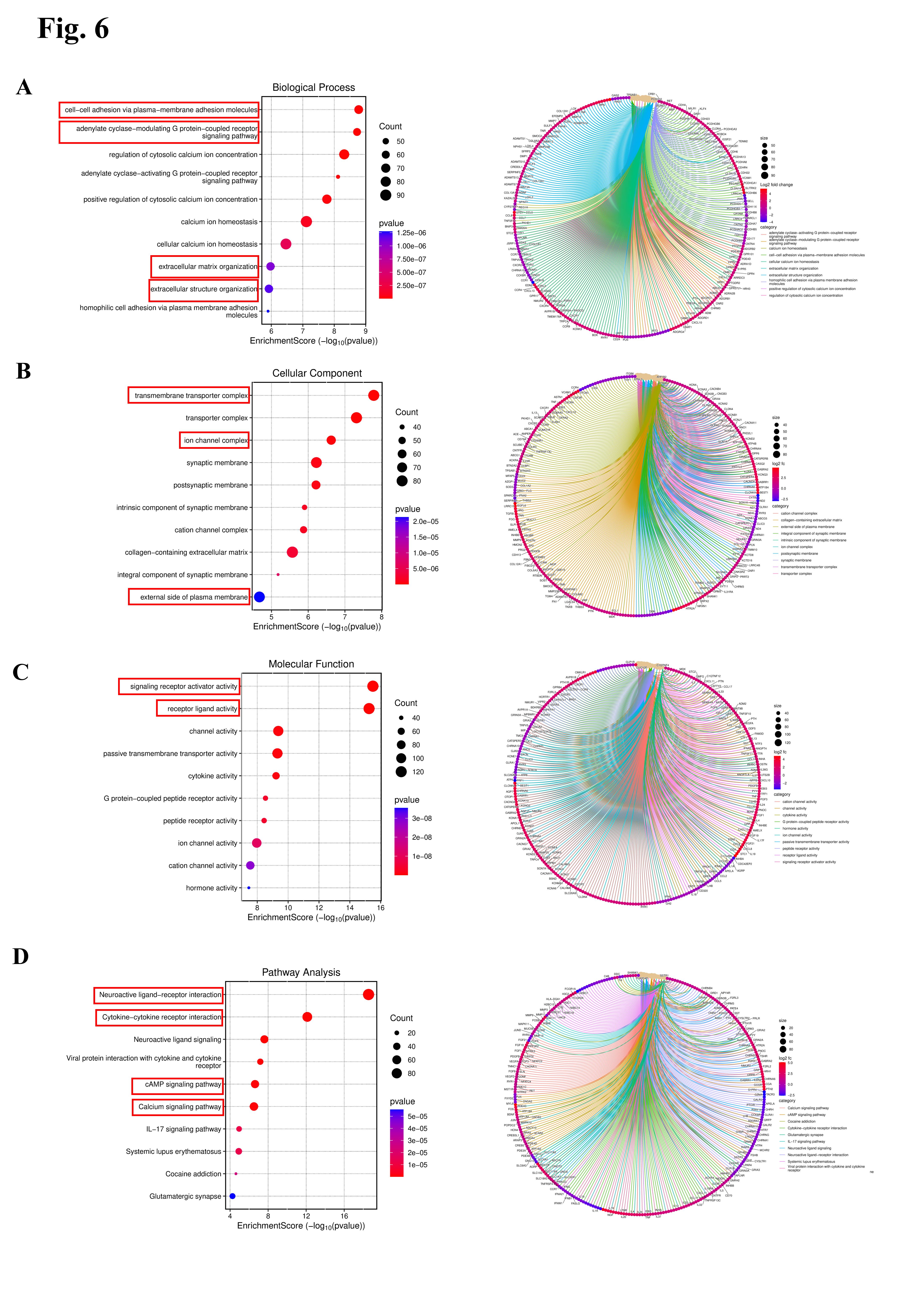
